# STCRpy: a software suite for T cell receptor structure parsing, interaction profiling and machine learning dataset preparation

**DOI:** 10.1101/2025.04.25.650667

**Authors:** Nele P. Quast, Charlotte M. Deane, Matthew I. J. Raybould

## Abstract

**Summary:** Computational methods to guide early-stage TCR drug discovery and TCR repertoire informatics currently under-utilise solved and predicted structure data. Here, we streamline use of these data through an open-source python package for high-throughput TCR structure handling and analysis (STCRpy), facilitating analyses such as TCR:peptide-MHC complex orientation calculation/scoring, root-mean-square-distance evaluation, interaction profiling, and machine learning dataset curation.

**Availability and implementation:** Freely available as a Python package at https://github.com/npqst/STCRpy.

## Introduction

T cell receptors (TCRs) direct the adaptive immune response by interacting with antigens, such as short peptide fragments, presented in mammalian cells by the Major Histocompatibility Complex (pMHC) (Birnbaum et al. [2014], Murphy and Casey [2017], van der Merwe and Dushek [2011]). They are gaining increasing attention, as a basis for the antigen targeting arm of biotherapeutics, especially for cancer, whether in soluble or cellular modalities (Dolgin [2022], Shafer et al. [2022]).

This coincides with the increased adoption of computational tools to streamline or reimagine the process of biotherapeutic molecule selection (Notin et al. [2024], Riley et al. [2016]). In the TCR field, this centres around unsupervised clustering or supervised prediction models that operate on the amino acid sequence and aim to assign the specificity of an expanded clone from a natural T-cell receptor repertoire (Hudson et al. [2023]).

These methods have so far shown promise in the context of thoroughly studied antigens, but are unable to generalise to unseen antigen contexts or more complex determinants of binding (Hudson et al. [2023], Meysman et al. [2022]).

A strategy to improve the generalisability of models is to consider 3D structural information in addition to the sequence (Robinson et al. [2021]). While a few structure-aware methods exist (Bradley [2023], Ghoreyshi and George [2023], Wang et al. [2024]), the overheads involved in accessing structural information and processing TCR:pMHC pose predictions are a roadblock to new entrants to the field. While numerous immunoglobulin-specific software packages exist for sequence-based analysis of TCRs (e.g. Dunbar and Deane [2016], Ye et al. [2013]), no such suite exists for annotating and processing TCR structure data, which are ever more abundant (Leem et al. [2018], Quast et al. [2025]).

One bespoke processing step is calculating the relative orientation of the TCR to the pMHC, a property linked to the ability of cell-surface TCRs to trigger downstream signalling (Zareie et al. [2021]) and to cross-reactivity and even auto-reactivity (Beringer et al. [2015], Gras et al. [2016]). Many general protein-protein complex prediction software packages generate poses of signalling TCRs that fall outside the ‘canonical’ angles consistent with this tenet of T cell biology. While methods have been proposed for calculating TCR geometries (Rudolph et al. [2006], Singh et al. [2020]), there is no standard method with an accessible python interface, limiting reproducibility.

Here, we present STCRpy, a python package to automate the processing and characterisation of solved and predicted TCR:pMHC complexes or *apo* TCRs.

## Implementation

STCRpy is pip-installable on Linux and Mac OSX operating systems, has been unit tested to assess the functionality of all modules (described below) and is provided with a python API and as a command line interface with full documentation.

### Parsing, annotation, and interaction analysis

TCR:pMHC complexes can be parsed from local PDB or MMCIF files, or pulled directly from STCRDab (Leem et al. [2018]) or the PDB (Berman et al. [2000]). Protein chains are first annotated using a modified version of ANARCI (Dunbar and Deane [2016]) able to classify MHC and MHC-like sequences as well as immune receptors (available as https://github.com/npqst/anarci-mhc). All TCRs are numbered and annotated by region according to the IMGT system/definitions (Lefranc et al. [2005]). TCRs and antigens are then paired by distance into separate ‘TCR’ objects, accounting for multiple TCR:pMHC copies per PDB file. As an extension of the BioPython PDB module (Cock et al. [2009]), STCRpy parses structures in a hierarchical manner (from ‘model’ to ‘chain’ to ‘residue’ to ‘atom’).

STCRpy processes each object with PLIP (Adasme et al. [2021]), characterising hydrogen bonds, salt bridges, and hydrophobic or aromatic interactions within and between all protein chains. The identified interactions can be extracted as a dataframe for further analysis (e.g. pose filtering, see Applications), or visualised as a heatmap or PyMOL (LLC and DeLano [2020]) session (Fig. 1a).

**Fig. 1.**
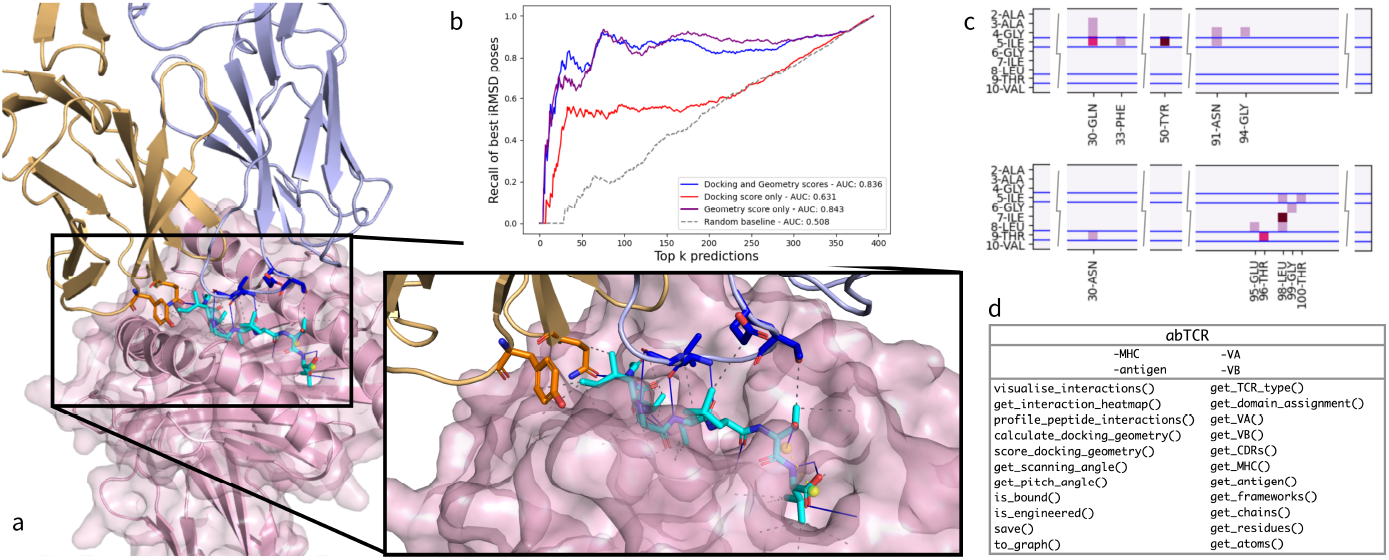
a) Visualisation of peptide interactions of *αβ*TCR in complex with pMHC (PDB ID: 6EQB, ‘bulged’ peptide conformation) generated with `tcr.visualise interactions()` from STCRpy. Base colours: *α* chain in orange, *β* chain in lilac, MHC in pink, peptide in cyan. Interface residues are highlighted as sticks using the standard heteroatom colour scheme, dashed lines depict interacting residues identified by PLIP (Adasme et al. [2021]). b) Recall of lowest interface RMSD (iRMSD) TCR:pMHC complex predictions from physics-based docking using energy-based docking scores (red, AUC of 0.631), STCRpy geometry score (purple, AUC of 0.843), and joint geometry and docking scores (blue, AUC of 0.836). Random baseline as grey dashes (AUC of 0.508). Using STCRpy docking geometry improves recall of best predictions, combining the docking score with the geometry score improves recall at *k <* 50. c) An example heatmap of interactions profiled between a peptide and the TCR *α* chain (top), and a peptide and the TCR *β* chain (bottom) generated by `tcr.get interaction heatmap()` from STCRpy. Peptide residues of interest (5-ILE and 9-THR) are highlighted with blue bars, pixel intensity intensity indicates the number of interactions. d) Table of TCR object-bound attributes and methods in STCRpy.

### TCR:pMHC Geometry Scoring and RMSD

We implement and define the geometry of a TCR relative to the MHC using three methods, two of which have previously been reported (Rudolph et al. [2006], Singh et al. [2020]), as well as an additional adaptation of the two methods, described in Appendix A, thereby unifying TCR:pMHC geometry evaluations within one software package. This enables consistent, comparable and reproducible calculations of TCR to pMHC geometry. Crucially, STCRpy distinguishes poses that fit the canonical angle/polarity from those that are reverse-canonical or non-canonical modes of engagement (Beringer et al. [2015], Gras et al. [2016], Zareie et al. [2021]), enabling systematic differentiation of poses compatible or incompatible with signalling during in-silico analyses.

We calculated the geometry of all complexes in STCRDab whose TCRs show evidence of downstream signalling, and supply these distributions in STCRpy, providing a means of scoring or filtering candidate docks (see Applications). Briefly, we parametrise the distribution of the scanning angle as a normal distribution, the pitch as a Gamma distribution, and the TCR to MHC distance as a mixture of Gaussians. Further, we calculate a score, *η*, as a linear combination of negative log-likelihood of a complex across the three distributions: *η* = Σ_*i*_ −*α*_*i*_*log*(*p*_*i*_(*x*_*T CR:pMHC*_)). We have reported the distribution parameters and their estimation in Appendix A, along with the implementation of the scoring function.

STCRpy also enables fast and reproducible calculations of RMSD both between TCR structures and across interfaces of TCR:pMHC complexes against a reference structure.

### Machine learning dataset preparation

STCRpy can parse both *apo* and bound TCR complexes into datasets of graphs using the popular pytorch-geometric and pytorch deep learning libraries (Fey and Lenssen [2019], Paszke et al. [2019]), facilitating the use of TCR structural information in machine learning. The generated graphs are compatible with the existing pytorch-geometric suite of graph neural network architectures, and can be integrated flexibly into training paradigms of bespoke neural networks. Furthermore, STCRpy enables users to flexibly assign labels, according to the neural network architecture and task they are interested in training. Alternative node selection, node featurisation, and edge featurisation methods can be configured, and bespoke implementations can be passed as arguments.

## Applications

### Evaluating predicted TCR:pMHC complex poses

We first demonstrate the utility of STCRpy by scoring and retrieving accurate docks from a pool of 2000 candidates using a TCR in complex with a KRAS-G12D antigen presented by HLA-A*11:01, deposited with PDB identifier 7PB2 (Poole et al. [2022]). As an *in-silico* complex prediction scenario, we predicted the TCR structure from sequence using TCRBuilder2+ (Quast et al. [2025]) and Alphafold-Multimer (Evans et al. [2022]). This resulted in nine candidate TCR structures, four from TCRBuilder2+ and five representative structures from Alphafold-Multimer, with which to initialise *in-silico* docking experiments. We report the RMSD of the TCR structure predictions, which can be easily and reproducibly calculated using STCRpy, in Appendix B Table 3.

We then used a physics-based approach (HADDOCK2.4, Van Zundert et al. [2016]) to dock the nine predicted structures and the original crystal structure of the 7PB2 TCR against the pMHC antigen. The docking simulations yielded 200 predicted TCR:pMHC complexes per TCR structure, resulting in a total of 2000 predicted TCR:pMHC complexes.

STCRpy enabled fast processing and evaluation of all predicted complexes *via* batch methods designed to handle large quantities of TCR structure data. For each pose, we used STCRpy to calculate the interface RMSD to the original crystal structure, quantify the geometry of the complexes and extract the energy-based docking scores from the HADDOCK simulations. Finally, we applied STCRpy’s geometry scoring functionality to each predicted pose.

The retrieval of plausible and accurate candidates would usually require manual inspection and evaluation, or reliance on docking scores which are relatively poor predictors of accuracy. Here, STCRpy enables consideration of both the energy-based docking score and a knowledge-based geometry score, which together correlate better with interface RMSD allowing retrieval of accurate TCR:pMHC complex predictions (Appendix B Fig. 6). Specifically, a simple linear regression of the docking and geometry score to the interface RMSD of the complex yields an AUPRC of 0.836 on the withheld validation set (Fig. 1b). Using exclusively geometry features results in a slightly higher AUC of 0.843, but worse retrieval at *k* < 50. The docking score feature alone underperforms relative to geometric scoring (AUC of 0.631), which underpins the difficulty of using general functions for evaluating docking methods.

### Characterising TCR:pMHC interface interactions

Identifying the interactions between the epitope and paratope of a TCR:pMHC complex is key to understanding the binding motif and provides insights into mutations that could impact binding affinity. As a case study we characterised the interactions between the HLA-A*02:01-AAGIGILTV antigen and two TCRs, MEL5 and *α*24*β*17, deposited in the PDB with the identifiers 6EQA and 6EQB respectively (Madura et al. [2019]). As Madura et. al. report, this antigen undergoes a shift in structural conformation from ‘stretched’ to ‘bulged’ upon TCR binding. We used the interaction profiling utility of STCRpy to compare the TCR:pMHC interfaces of both peptide states, showing that additional hydrophobic contacts between the TCR alpha chain and the peptide emerge in the immunogenic state (Appendix C Table 4). For the full analysis, see Appendix C. The interactions, which are generated using PLIP (Adasme et al. [2021]), can be visualised as an annotated heatmap (Fig. 1c, Appendix C Fig. 7) or in PyMOL (LLC and DeLano [2020], Appendix C Fig. 8).

### Machine learning TCR property prediction

STCRpy’s `TCRGraphDataset` constructor converts TCR structures into graphs compatible with graph neural networks (GNNs). The graphs can be configured; by default each amino acid is defined as a node and edges correspond to distances between amino acids. We validated our graph datasets by training an equivariant graph neural network (EGNN) (Satorras et al. [2021]) on region annotation of a point cloud, whereby the network assigns a region label, TCR*α*, TCR*β*, peptide, or MHC, to each amino acid, based on the structure of the complex alone (see Appendix D). The attained node annotation accuracies on the validation set substantially outperform the random baseline (Appendix D Table 6), demonstrating that EGNNs learn the underlying geometry of TCR:pMHC complexes from the STCRpy graph inputs.

## Conclusion

We have illustrated STCRpy’s utilities (Fig. 1d) with three indicative workflows. Firstly, we have showed that STCRpy can be used to evaluate predicted TCR:pMHC complexes for TCR-likeness using a geometry score. In this case we generated poses using a physics-based approach, but STCRpy could also be applied to the outputs of machine learning models such as AlphaFold3 (Abramson et al. [2024]). Secondly, we have demonstrated that STCRpy can be used to profile interactions between TCRs and peptides, specifically demonstrating a case where sequence analysis alone would not suffice as the interactions arise from a change in conformation. These interaction profiling methods could be exploited to generate further TCR-likeness metrics (Raybould et al. [2023]) beyond the geometry score used in the first case study. Finally, we have shown that STCRpy can be used to convert TCR structures to graph datasets for deep learning pipelines, removing barriers to the use of predicted structural features in TCR:pMHC specificity classifiers.

Overall, STCRpy offers a simple route toward analysing the broad diversity of structural data on TCRs and their complexes emerging from repertoire studies and TCR-based drug discovery. We have released the STCRpy codebase as an extendible open source project, and encourage community engagement and contributions as the field evolves.

## Author contributions statement

N.Q., C.D. and M.R. conceived the work. N.Q. conducted the work. C.D. and M.R. supervised the work. N.Q., C.D. and M.R. analysed the results. N.Q. wrote the manuscript and C.D. and M.R. reviewed and edited the manuscript.

## Acknowledgments

This work is supported by the Engineering and Physical Sciences Research Council (EPSRC, grant EP/S024093/1) and Immunocore Ltd. The authors thank and acknowledge James Dunbar, Jinwoo Leem and Catherine Wong for their contributions to early versions of the TCR parsing modules through their work on ANARCI and the SAbDab and STCRDab databases from 2013-2020.

## Appendix A: Geometry calculation and scoring of TCR:pMHC complexes

### Geometry calculation

Three methods for calculating geometric metrics of TCR:pMHC complexes are implemented in STCRpy. All three methods calculate the scanning angle of TCR to pMHC slightly differently and we have described and characterised these differences here. The third method, which is a combination of both methods we have implemented in STCRpy, is also used to score docking geometry.

### Rudolph et al

This method for calculating the scanning angle of a TCR bound to MHC (Rudolph et al. [2006]) is implemented in STCRpy as follows:

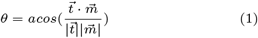

Where *θ* is the scanning angle of the TCR over the MHC, 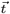 is the vector pointing from the TCR*α* chain to the TCR*β* and 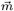 is the vector pointing through the MHC. More specifically, the TCR vector is defined as the unit-normed vector pointing from the mean, *i*.*e*. centroid, of the *α* chain cysteine residues’ sulphur atom to the mean of the *β* chain cysteine residues’ sulphur atom, where in each chain the cysteine residues comprise the conserved variable domain disulfide bridge (between positions 23 and 104 using IMGT numbering, Lefranc et al. [2005]):

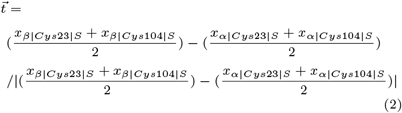

The MHC vector is defined as the unit-normed vector pointing through the MHC. This vector is determined as the principal component of the singular value decomposition (SVD) of the point cloud of coordinates of the MHC helices forming the peptide cleft, *X*.

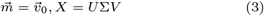

where 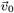 is the first component of the *V* matrix of the SVD. Intuitively, *v*_0_ explains the most variance in the coordinates *X* in 3D Euclidian space. The point cloud *X* ∈ ℝ^3×*N*^ is defined as the coordinates of the *C*_*α*_ of the MHC helices, translated so the centre of mass is at the origin:

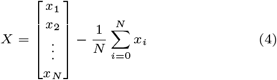

The *N* residues comprising the helices are selected by residue numbering in Table 1. We use the `numpy.linalg.svd` algorithm to solve the decomposition. Finally, the sign of the vector, the direction of which is stochastic, is aligned to have the same direction as the approximate MHC vector, 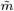, pointing from the N to the C terminus of the *α*_1_ helix:

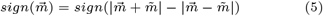

In STCRpy, this method for calculating the scanning angle angle can be called as follows: tcr.calculate geometry(mode=‘rudolph’).

### Centre of mass (Singh et al. [2020])

An advantage of the Rudolph *et al*. method is that it does not require the definition of a global coordinate frame – both the TCR vector and the MHC vector are calculated within the same frame, making the angle between them invariant to rotations. However, this limits the evaluation of the TCR:pMHC geometry to metrics invariant to rotations and translations. It can also lead to spurious edge cases as the scanning angle *θ* is invariant to rotations of the TCR vector about the axis of the MHC vector. While this is not necessarily a limitation for well-behaved crystal structure of TCR:pMHC complexes, *in-silico* predictions often contain noisy and non-canonical predictions that this method fails to distinguish by scanning angle alone. An extreme example would be a TCR docked to an MHC from the side at a 90^°^ angle, which would be annotated with a canonical value of *θ* if calculated using the Rudolph *et al*. method.

**Appendix A Table 1.** MHC residue number ranges included in point cloud coordinates when calculating MHC vector 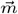 as in Eq. 3&4. Different numbering applies for MHC Class I and Class II. In Class I both helices are encoded by the same chain, whereas for Class II the alpha helices are on different chains. The two ranges for MHC Class II *α*_2_ exclude the ‘kinked’ region of the second helix.

| | $\alpha_1$ | $\alpha_2$ |
| --- | --- | --- |
| MHC Class I | (50-87) | (140-177) |
| MHC Class II | (50-88) | (54-65) & (67-91) |

As an alternative, Singh *et al*. proposed to align TCR:pMHC complexes to MHC reference structures through which a Cartesian coordinate frame has been defined (Singh et al. [2020]). Specifically, the origin is defined to be at the centre of the peptide cleft, with the x-axis cutting across the cleft pointing from MHC helix *α*_1_ to *α*_2_, the y-axis running along the peptide cleft, and the z-axis pointing upwards toward the TCR interface. To calculate the TCR:pMHC complex geometry, first the TCR:pMHC complex is aligned by MHC to the correct reference structure (MHC Class I or Class II). After this alignment is complete, the TCR:pMHC sits in the predetermined global coordinate system, and absolute metrics, which are subject to translations and rotation, can be evaluated and compared across TCR:pMHC complexes.

Within this coordinate frame, we follow Singh *et al*.’s definition of the scanning angle *θ*, pitch *ψ*, and z-distance, to define the geometry of the TCR:pMHC complex. The scanning angle is defined as the angle between the projection 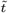 of the TCR vector 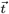 onto the xy-plane and the y-axis:

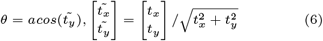

The pitch angle *ψ* of the TCR:pMHC complex is defined as the angle between the TCR vector 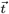 and the z-axis. Intuitively this describes the tilt of the TCR.

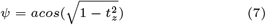

**Appendix A Fig. 2.**
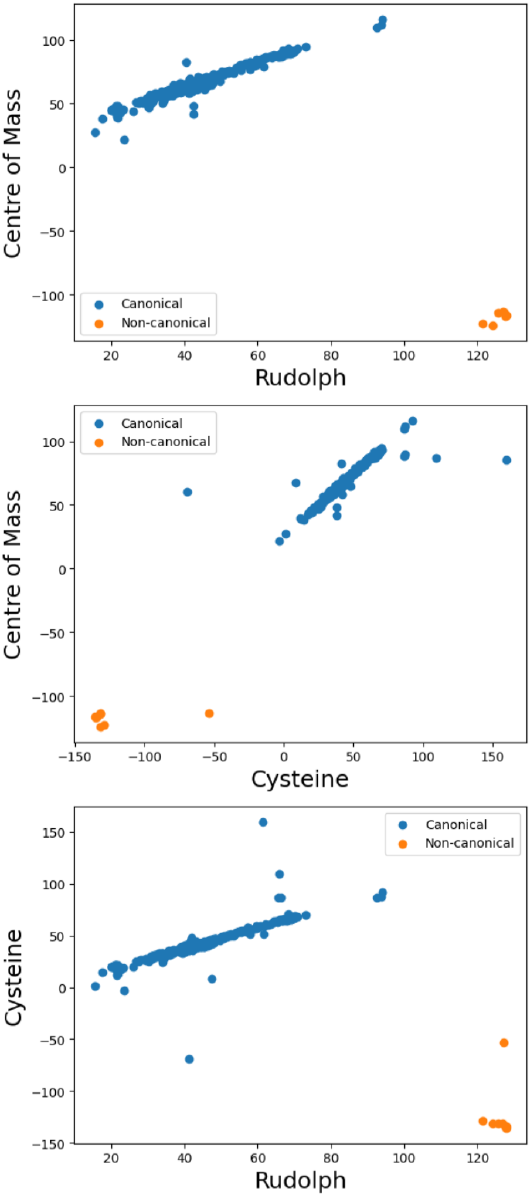
Comparison of MHC Class I TCR:pMHC scanning angles using three methods implemented in STCRpy. The methods produce linearly related scanning angles, with the exception of outliers, but the scanning angle ranges differ across methods. Non-canonical binding modes (orange) are readily separable across all three methods, with the exception of an outlier with additional engineered cysteine residues.

**Appendix A Fig. 3.**
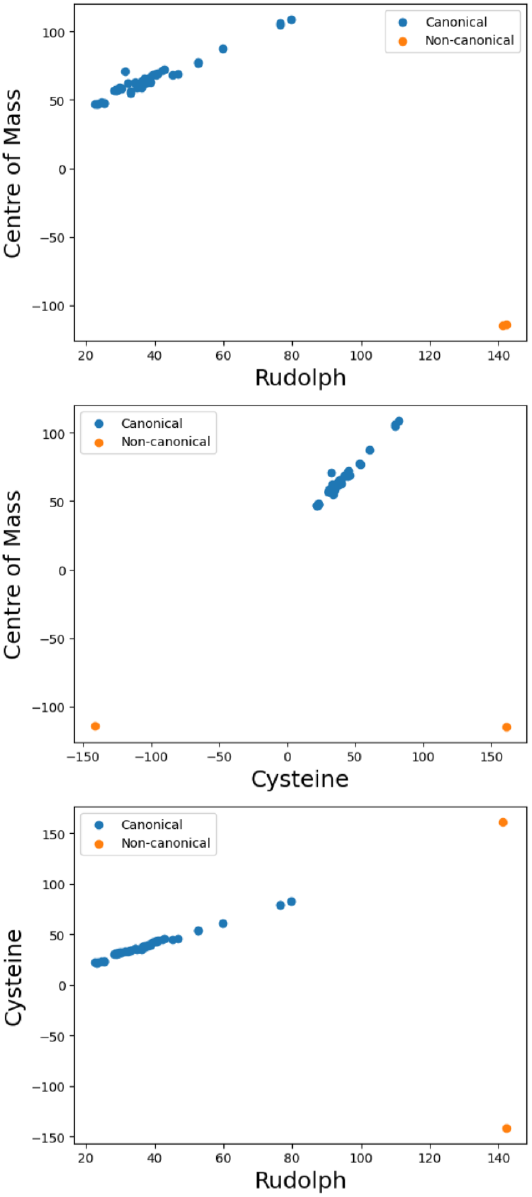
Comparison of MHC Class II TCR:pMHC scanning angles using three methods implemented in STCRpy. The methods produce linearly related scanning angles, with the exception of outliers, but the scanning angle ranges differ across methods. Non-canonical binding modes (orange) are readily separable across all three methods.

The definition of the TCR vector differs from Rudolph et.al. - instead of defining the angle pointing from the *α* chain to the *β* chain via the cysteine residues’ centroid, Singh *et al*. calculate the centre of mass of a subset of each chain, and uses the vector between these points:

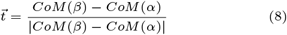

The residue subset used for each TCR chain is provided by a reference file included in STCRpy for each TCR chain type, *α, β, γ*, and *δ*.

In STCRpy, this method for calculating the TCR:pMHC geometry can be called as follows: tcr.calculate geometry(mode=‘com’)

### Combined cysteine and centre of mass method

We have also implemented a new, combined way of calculating TCR:pMHC geometric features, which takes advantage of Singh *et al*.’s alignment to a global coordinate frame while avoiding the dependence on a reference residue subset for calculating the TCR centre of mass. First, we align the TCR:pMHC to a reference MHC, but then, we calculate the TCR vector using the cysteine residue centroids as in Eq. 2. The remainder of the TCR geometry is then calculated following Eqs. 6 & 7. In STCRpy, this method for calculating the TCR:pMHC geometry can be called as follows:

tcr.calculate geometry(mode=‘cys’)

**Appendix A Table 2.** Parameters of probability distributions used to calculate STCRy geometry score using maximum likelihood estimation on STCRDab data.

|  | Distribution type | Parameters |
| --- | --- | --- |
| $\theta$ | Normal | $\mu = 67.9, \sigma^2 = 202.1$ |
| $\psi$ | Gamma | $\alpha = 1.20, \theta^{-1} = 5.78,$<br>$x = 0.0365$ |
| $z - dist$ | Gaussian Mixture | $N = 2,$<br>$\mu = [27.05, 28.62],$<br>$\sigma^2 = [0.468, 0.908],$<br>$w = [0.434, 0.566]$ |

An advantage of the global coordinate frame is the intuitive identification of canonical vs. non-canoncial TCR:pMHC binding modes: given the coordinate frame, any TCR vector with a negative x-component, ie. the vector from the TCR *α* chain to the TCR *β* chain points ‘upwards’ at the *α*_1_ helix, has a canonical binding geometry. This holds true for both methods “centre of mass”, and “cysteine” which align the TCR:pMHC to a global reference frame. Using the method proposed by Rudolph *et al*. it is not possible to use absolute components of the TCR or MHC vector, since these are subject to changes under rotation and translation. However, empirically, we find that all scanning angles *θ >* 120^°^ correspond to a non-canonical, reverse binding TCR:pMHC geometry. In STCRpy, the polarity can also be embedded into the scanning angle, by multiplying the angle by −1 for reverse polarity. In Fig.2&3 this has been applied to the ‘cysteine’ and ‘centre of mass’ methods. To enable comparisons across these geometry calculations and to confirm their self-consistency, we characterised the methods using TCR:pMHC crystal structures from STCRDab (Appendix A Figs. 2 & 3). We observe that the relationship between scanning angles across all methods is linear, as expected, and that non-canonical binding modes are readily separable.

### Geometry scoring

We are able to score predicted TCR:pMHC complexes by evaluating the likelihood that they arose from the known TCR:pMHC distribution. We featurise the TCR:pMHC complexes by geometric features: the scanning angle *θ*, the pitch angle *ψ*, and the z-component of the TCR centre of mass, which we calculate using the cysteine method described above. We calculate the distribution of these metrics over TCR:pMHC complexes in STCRDab, and fit simple parametric distributions to the data using maximum likelihood estimation. We then evaluate the likelihood of any query TCR:pMHC complex arising from these ‘TCR-like’ distributions:

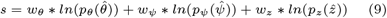

Where 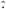 is the geometric feature of the predicted query complex. The weights *w* can be adjusted but are set to *w*_*θ*_ = *w*_*ψ*_ = *w*_*z*_ = 1 by default. We approximate *θ* with a normal distribution, *ψ* with a gamma distribtuion, and z-distance with a mixture of Gaussians. We fit the distributions using the popular scikit-learn and scipy implementations. We have reported the optimal parameters in Appendix A Table 2 and visualised the fit in Appendix A Fig. 4.

**Appendix A Fig. 4.**
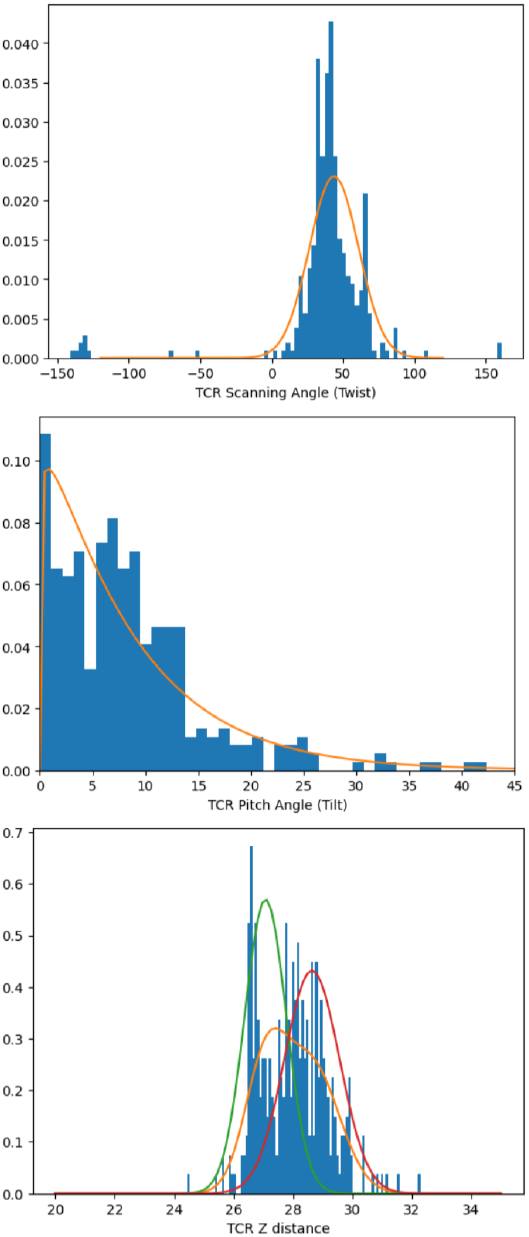
Parametric probability distributions fit to STCRDab MHC Class I data. Scanning angle *θ* is fit with a normal distribution, pitch angle *ψ* with a gamma distribution, and z-distance with a mixture of Gaussians, n=2. Blue histogram bars are STCRDab data, orange line is the maximum likelihood estimate (MLE) fit of the parametric distribution. Red and green lines are the distributions of the independent components of the Gaussian Mixture.

## Appendix B Evaluating predicted TCR:pMHC complexes

We used physics-based docking to generate 2000 TCR to KRAS-G12D HLA-A*11:01 complexes and analysed these predicted structures with STCRpy.

### Results

We present the RMSD of the structure predictions used for docking in Table 3.

Plotting the scanning angle against the HADDOCK energy score demonstrates that when the TCR structure is not known, a wide range of potential TCR:pMHC complex poses are sampled, including some which are not conducive to TCR signalling due to a non-canonical binding mode (Beringer et al. [2015], Gras et al. [2016], Zareie et al. [2021]), (Appendix B Fig. 5).

**Appendix B Fig. 5.**
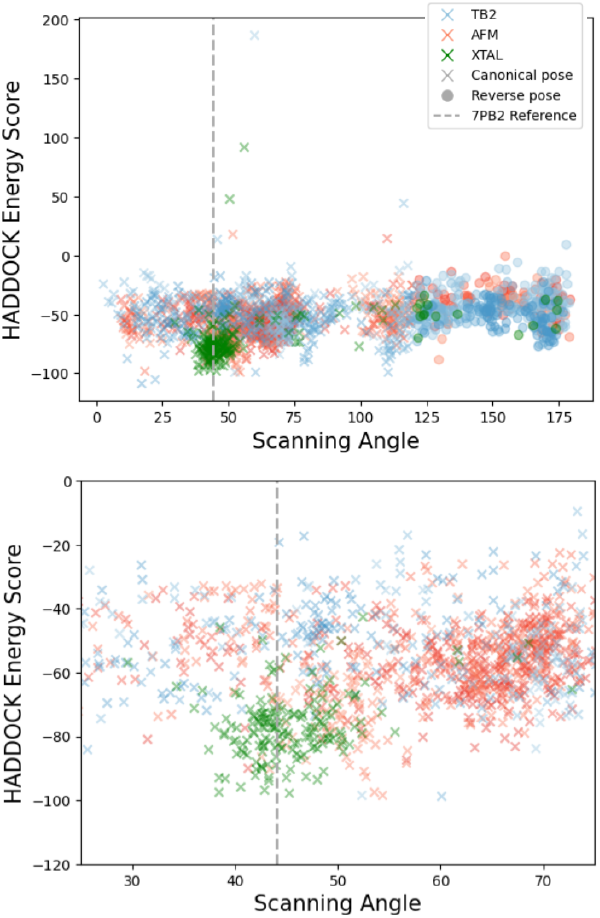
Haddock energy score of TCR:pMHC docks against scanning angle of the TCR to MHC of 7PB2 case study. Grey dashed line is the true scanning angle of the crystal structure. AFM: docks using Alphafold-Multimer TCR structure predictions, TB2: docks using TCRBuilder2+ TCR structure predictions, XTAL: docks using the experimental crystal structure of the TCR. x and o markers represent canonical and non-canonical scanning angles, respectively. When the TCR structure is predicted, rather than known, the range of predicted TCR:pMHC scanning angles increases substantially, and the docking score is approximately uniform across angles, making the retrieval of accurate or ‘TCR-like’ poses based on the docking score impossible.

However, when considering the interface RMSD plotted against the HADDOCK energy score and geometry scores it becomes possible to identify valid poses more readily, with lower interface RMSD clearly correlated to lower geometry scores (Appendix B Fig. 6).

**Appendix B Fig. 6.**
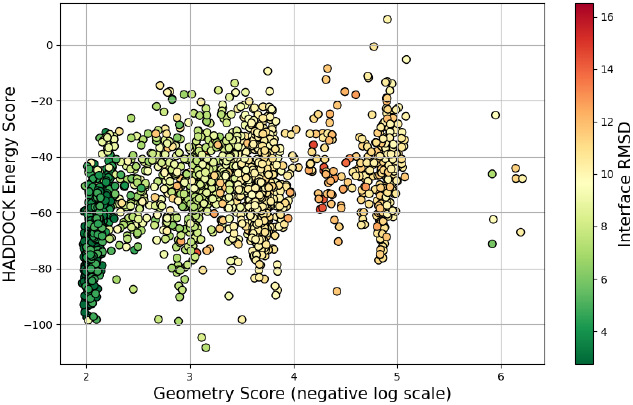
Haddock energy score *vs*. the negative natural logarithm of the STCRpy geometry score of TCR:pMHC docks, coloured by interface RMSD [Å] to the 7PB2 ground truth crystal structure. Geometry score is better correlated with low interface RMSD than the docking score.

### Methods

We predicted the TCR structure of the 7PB2 TCR using TCRBuilder2+ (Quast et al. [2025]) and Alphafold-Multimer (Evans et al. [2022]), retrieving four and 25 predictions from each model respectively. For the Alphafold-Multimer structures we clustered the structures into distinct conformations with greedy clustering and a 1Å threshold, yielding 5 distinct predictions. We used these five predictions in addition to the four TCRBuilder2+ predictions for docking.

We prepared the structures for docking by separating the antigen from the TCR of the 7PB2 crystal structure. We then used the reformatting functions implemented in SCTRpy to renumber the structures and assign all residues to single chains, making them compatible with the docking software, HADDOCK2.4 (Van Zundert et al. [2016]). We defined the CDR loops of the TCR and the peptide residues as ‘active’ residues in the simulation; HLA residues within 6Å of the peptide were defined as ‘passive’ residues. The remaining parameters of the HADDOCK simulations were left as default. We docked the nine structure predictions and the TCR crystal structure to the antigen structure, yielding 2000 predicted TCR:pMHC complexes in total.

After the docking simulations were complete we used the STCRpy reformatting methods to restore the numbering of the TCR:pMHC complexes and parsed docking metrics, such as the energy-based docking score from the simulation. We then calculated the interface RMSD to the crystal structure of the complex, the scanning angle, pitch angle, and z-distance (using the ‘centre-of-mass’ method) of the TCR relative to the antigen, as well as the geometry scores of each predicted complex. We further profiled the peptide interactions of every complex. Using STCRpy this analysis reduces to a single line, which saves the retrieved metrics to CSV files at a specified location:

~~~
1      stcrpy. tcr_methods. tcr_batch_operations .
           analyse_tcrs (
2                  tcr_complex_files,
3                  save_dir=“.”)
~~~

To retrieve a docking complex from the 2000 candidates, we rejected any TCR:pMHC complexes where no interactions were profiled between the peptide and the TCR. We then rank the remaining docks by their geometry score and select the candidate with the lowest score.

To investigate the retrieval power of the docking scores and geometry scores, we used a simple linear regression model to predict the interface RMSD, `iRMSD`. The linear regression was fit on 80% of the predicted TCR:pMHC complexes. We fit three models, one with both the energy based docking term, *x*_*D*_, and the natural logarithm of the negative geometry score,

**Appendix B Table 3.** RMSD of TCR structure predictions used for the physics based docking case study evaluated against the 7PB2 crystal structure. TB2+: TCRBuilder2+(Quast et al. [2025]), AFM: Alphafold-Multimer (Evans et al. [2022]). ‘-X’ denotes the internal rank of the structure prediction assigned by the prediction model. RMSD for the whole TCR and across regions can easily be calculated using: `stcrpy.tcr metrics.RMSD().calculate rmsd(prediction, reference)`, where `prediction` and `reference` are ‘TCR’ objects.

|  | TB2+ - 0 | TB2+ - 1 | TB2+ - 2 | TB2+ - 3 | AFM - 0 | AFM - 4 | AFM - 6 | AFM - 12 | AFM - 15 |
| --- | --- | --- | --- | --- | --- | --- | --- | --- | --- |
| A | 0.89 | 1.07 | 1.20 | 1.36 | 1.35 | 1.23 | 0.89 | 0.87 | 0.89 |
| CDRA1 | 1.20 | 1.78 | 1.12 | 1.13 | 0.72 | 0.80 | 0.63 | 0.47 | 0.46 |
| CDRA2 | 1.35 | 0.84 | 1.40 | 1.26 | 1.88 | 1.90 | 1.81 | 1.72 | 1.77 |
| CDRA3 | 1.36 | 2.01 | 2.31 | 2.85 | 3.42 | 2.99 | 1.80 | 1.81 | 1.87 |
| FWA | 0.69 | 0.77 | 0.90 | 0.98 | 0.49 | 0.50 | 0.47 | 0.45 | 0.45 |
| B | 0.99 | 1.20 | 0.97 | 0.81 | 1.22 | 1.19 | 1.02 | 1.31 | 1.32 |
| CDRB1 | 0.32 | 0.45 | 0.53 | 0.27 | 0.49 | 0.47 | 0.42 | 0.38 | 0.44 |
| CDRB2 | 0.88 | 1.16 | 1.14 | 0.70 | 0.72 | 0.69 | 0.61 | 0.78 | 0.84 |
| CDRB3 | 2.38 | 2.93 | 2.22 | 1.75 | 2.05 | 1.59 | 0.95 | 2.33 | 2.42 |
| FWB | 0.58 | 0.64 | 0.58 | 0.58 | 1.10 | 1.17 | 1.07 | 1.14 | 1.12 |

*ln*(−*x*_*g*_). We then ranked the predicted interface RMSD of the remaining 20% of complexes, and calculated the recall of the top *k* predictions as 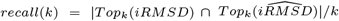, where 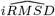 is the predicted interface RMSD. We report the AUC of the recall curve calculated as 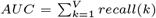, where *V* is the size of the validation set.

**Appendix C Fig. 7.**
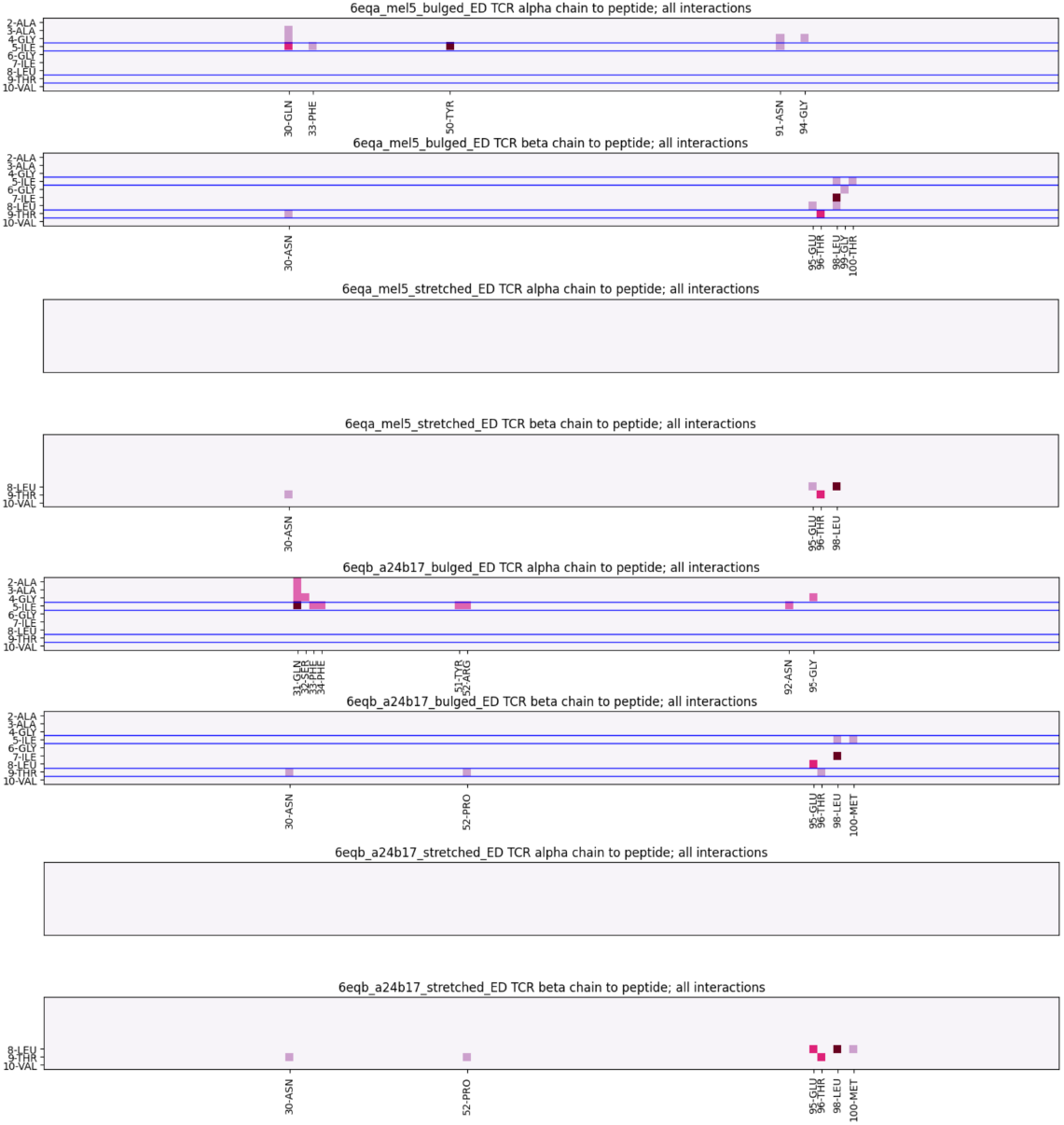
STCRpy generated TCR to peptide interaction heatmaps of 6EQA and 6EQB complexes in both the ‘bulged’ and ‘stretched’ peptide conformations. Pixel intensity indicates the number of profiled interactions. As reported by Madura et al. [2019], additional interactions emerge between the peptide and the TCR in the bulged peptide conformation. Specifically, interaction between the CDR*α* chains and the proximal peptide residues 2-ALA through to 5-ILE emerge.

## Appendix C Interaction profiling of 6EQA and 6EQB

### Results

Using STCRpy, we have analysed the ‘bulged’ and ‘stretched’ conformations of the 6EQA and 6EQB TCR:pMHC complexes’ interaction profiles. We show the interaction heatmaps in Appendix Fig. 7, as well as the visualised interactions of the structures in Appendix Fig. 8. We have quantified the interactions profiles in Appendix Table 4. As reported in the original paper (Madura et al. [2019]), the TCR-induced ‘bulged’ state of the peptide results in additional interactions between the TCR and antigen, which are expected to facilitate binding and signalling of the TCR.

**Appendix C Table 4.** Interactions between the TCR (MEL5 and *α*24*β*17) and the peptide in both stretched and bulged conformations as profiled by *STCRpy* using PLIP (Adasme et al. [2021]). More interactions between the peptide and TCR emerge in the bulged peptide conformation.

| | MEL5 | | $\alpha 24\beta 17$ | |
| --- | --- | --- | --- | --- |
|  | Bulged | Stretched | Bulged | Stretched |
| hbond | 4 | 0 | 5 | 0 |
| hydrophobic | 1 | 1 | 2 | 1 |
| saltbridge | 0 | 0 | 0 | 0 |

### Methods

TCR:pMHC crystal structures deposited in the PDB with the identifiers 6EQA (MEL5) and 6EQB (*α*24*β*17) with the stretched and bulged conformations recorded as alternative coordinates with the PDB record. After separating the conformations, we use the following STCRpy functions to profile and visualise the interactions:

~~~
1      *# load TCR structure and profile interactions*
2      tcr_complex = stcrpy. load_TCR (PDB_FILE)
3      interactions_dataframe =
4          tcr_complex. profile_peptide_interactions ()
5      tcr_complex. get_interaction_heatmap ()
6      tcr_complex. visualise_interactions ()
~~~

To demonstrate the flexibility of STCRpy we adjust the thresholds of the interaction identifier; the full interface can be found in the package documentation. In this study we use the parameter configurations specified in Appendix C Table 5.

**Appendix C Table 5.** Parameter configurations used for profiling interactions. All units in Ångström. *d*_*BS*_ - maximum distance from the peptide at which to include binding site residues; 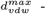 maximum distance at which to detect hydrophobic contacts; 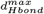 maximum distance between hydrogen bond donor and acceptor. All other interaction parameters were set to their default values, for more details see the *STCRpy* documentation.

| | $d_{BS}^{max}$ | $d_{vdw}^{max}$ | $d_{Hbond}^{max}$ |
| --- | --- | --- | --- |
| Nr. detected interactions (Tab. 4) | 7.5 | 4.0 | 4.1 |
| Interaction heatmap (Fig. 7) | 10. | 7.5 | 6.0 |
| 3D visualisation (Fig. 8) | 10. | 5. | 4.5 |

## Appendix D Graph Machine Learning with STCRpy

STCRpy can generate graph objects and datasets compatible with graph neural networks in deep learning pipelines. We validated our graph datasets by training the popular equivariant graph neural network (EGNN) architecture on a region annotation task, whereby the model classifies amino acids in the graph as belonging to the TCR*α*, TCR*β*, peptide, or MHC chains.

### Results

We report the node classification accuracies for region annotation in Appendix D Table 6. The EGNN attains good accuracies considering the size of the network, particularly on peptide and MHC residue annotation, which are relatively undersampled in the dataset. As the EGNN retains equivariance by considering the relative distances between coordinates to update node features, the ability of a small network to recapture the regions implies that the graph representation of TCR:pMHC complexes encodes structural information specific to these complexes. For context, the random baseline is achieved by sampling node classes randomly from a biased distribution, where the bias is determined by the relative frequencies of node labels.

**Appendix C Fig. 8.**
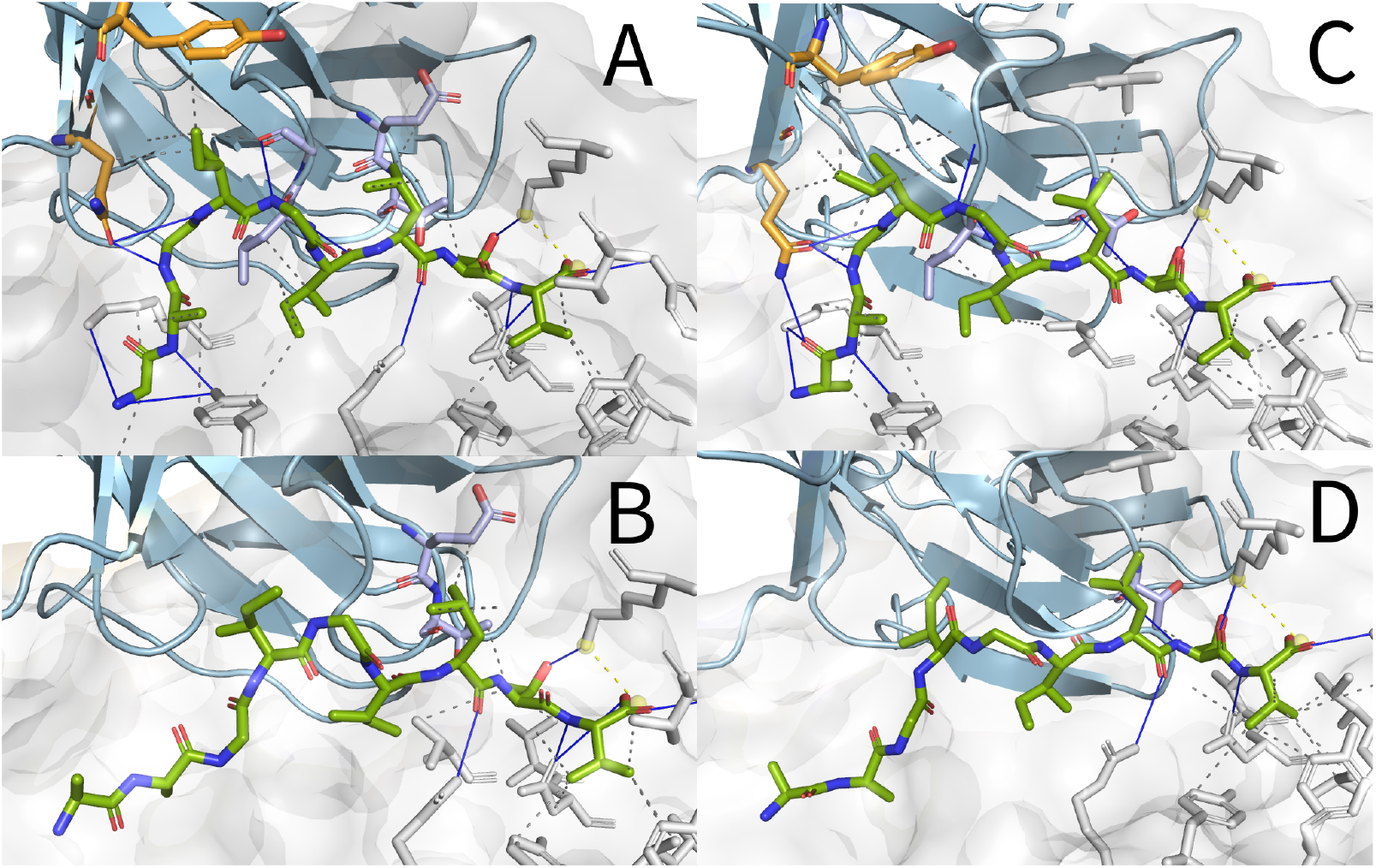
Peptide interactions of ‘bulged’ (A & C) and ‘stretched’ (B & D) peptide conformations visualised using STCRpy interface for PLIP (Adasme et al. [2021]) and PyMOL (LLC and DeLano [2020]). Peptide in green, MHC in grey, TCR in blue. Interacting TCR *α* and *β* chain residues in orange and lilac respectively.

**Appendix D Table 6.** Node classification accuracy on validation set for each TCR:pMHC region.

| Region | EGNN | Random Baseline |
| --- | --- | --- |
| TCR $\alpha$ | 94.68% | 37.43% |
| TCR $\beta$ | 96.30% | 47.87% |
| Peptide | 84.30% | 12.73% |
| MHC | 82.56% | 2.08% |

### Methods

We use the STCRpy graph dataset generator to re-format TCR structures from STCRDab (Leem et al. [2018]) to `torch geometric` compatible graphs. Node and edge featurisation is flexible, here we embed the residue and atom type as a one-hot encoding, and pass the atom coordinates as node associated positions. We opt to define a fully-connected graph with binary edges, since the EGNN calculates the distance and selects a maximum number of nodes with which to update the nearest neighbours internally. To reduce the graph size we opt to include only the carbon alpha of each residue, and only include MHC residues within 15 Ångström of any TCR residues. The final graph representation is akin to a point cloud of amino acids coordinates, with edges corresponding to their pairwise distances. We labelled each node by region (TCR *α*, TCR *β*, peptide, or MHC) and used these annotations as the classification target. We use a random 3:1 split for training and validation, resulting in 319 and 107 training and validation structures, respectively.

We built an equivariant graph neural network using the `egnn` package (Satorras et al. [2021]). Specifically, we define a linear node embedding layer, which projects the one-hot node encodings into 32 dimensions. We then pass the embeddings and the residue coordinates through three equivariant graph network layers, along with the fully-connected adjacency matrix. The graph layers constitute a message passing network that achieves SE(3) equivariance by considering only the relative distance between nodes rather than explicit coordinates, for details, refer to Satorras et al. [2021]. We select a maximum of 16 neighbouring nodes to be included in each node update, and use an internal node embedding dimension of 32. As we are performing node classification we do not update the coordinate positions as the graph proceeds through the network. We pass the final node embeddings through an output network consisting of two linear layers with an intermediate ReLU nonlinearity to obtain the class probabilities as logits. We trained the network using weighted cross-entropy loss and the Adam optimiser without weight decay. We defined the cross-entropy weights as the inverted frequency of classes in a random sample: *w* = [2.6471, 2.0886, 7.9839, 55.0000], thereby increasing the weight of the under-represented classes. We trained the network to convergence on the validation set.

